# Specialized Bacteroidetes dominate the Arctic Ocean during marine spring blooms

**DOI:** 10.1101/2023.09.26.559482

**Authors:** Álvaro Redondo-Río, Christopher J. Mundy, Javier Tamames, Carlos Pedrós-Alió

## Abstract

A metagenomic time series from Arctic seawater was obtained from the Canadian region of Dease Strait, to analyse the changes in bacterioplankton caused by the phytoplankton bloom that recurrently occurs in summer. This dataset documents the growth of bacterial clades specialised in the metabolism of plysaccharides, such as Bacteroidetes, along with the phytoplackton. These specialised taxa quickly displaced the microbial clades that dominate nutrient-poor waters during early spring, such as Archaea, Alpha- and Gammaproteobacteria. At the functional level, phyla Bacteroidetes, Planctomycetes and Verrucomicrobia showed higher contents of polysaccharide-degradation functions. Glycoside hydrolases revealed that the Bacteroidetes community shifted towards species with higher polysaccharide-degrading capabilities, targeting algal polysaccharides in summer. Regarding transporters, Bacteroidetes dominated SusC-TonB transporters and had an exclusive family of glycoside-binding proteins (SusD). These proteins were used to identify polysaccharide-utilisation loci that clustered transporters and polysaccharide-active enzymes, showing a higher level of specialisation towards polysaccharide use. Put together, all these genomic features point to the genetic adaptations that promote the dominance of Bacteroidetes during phytoplankton blooms.

## Introduction

Marine primary production is a main contributor to global carbon fixation, accounting for about half the global photosynthetic carbon assimilation (1). Recurrent phytoplankton blooms are an example of the capability of marine organisms to fix substantial amounts of carbon into organic matter. In these events, marine photosynthetic microorganisms flourish in coastal areas during spring or summer, driven by the rise in temperature, the increase in solar irradiation and the build-up of nutrients during winter.

During these blooms, the lysis of microalgae by predators and viruses causes an increase in dissolved organic carbon (DOC), including diatom-synthesised polysaccharides. This DOC enrichment provides a suitable substrate for the proliferation of heterotrophic bacteria, which in turn remineralise the bulk of the fixed carbon. Some polysaccharides and particles that are resistant to bacterial hydrolysis undergo sedimentation into the deep ocean, acting as a carbon sink. Therefore, these events are crucial for comprehending global carbon cycling.

Phytoplankton blooms and the consequent bacterioplankton blooms have been thoroughly studied in most temperate regions (2–5) using single-cell labelling (BIC-FISH or CARD-FISH) and metagenomics. Since the main bacterial substrates for during these events are polysaccharides, some studies have paid especial attention to the use that bacterioplankton make of these polymers (5). Polysaccharides are synthesised by microalgae as cell wall constituents, carbon storage and extracellular mucilages. It has been estimated that laminarin alone, the primary storage polysaccharide in diatoms, represents 5-15 Gt of biomass produced annually (6). However, most planktonic heteroglycans are still undescribed, with only some analyses revealing a wide variety of monomers and bonds, and a noticeable diversity among phytoplankton species (7).

Studies in polar regions have proven that water below ice receives enough solar radiation to sustain growth of ice-associated diatoms in the Arctic (8–10) and recent expeditions have observed microalgae blooms in the Southern Ocean under Antarctic ice (11).

Planktonic diatoms are mainly responsible for spring-summer blooms in the Arctic Ocean, with a dominance of, among others, *Nitzschia frigida* and *Attheya ssp.*, belonging to clade Bacillariophyta, although dinoflagellates (Dinophyceae), *Phaeocystis* (Haptophyceae) and picoeucaryotes are also present in this environment and contribute to the phytoplankton blooms (12).

Bacterial communities in both polar regions have been studied through 16S-rRNA analyses in the context of algal blooms (13–15), but only recently metagenomic has been applied to Arctic samples (16,17). Despite the existing research, a metagenomic time series covering a phytoplankton bloom is still lacking in the Arctic ecosystem, and so does a closer examination of polysaccharide utilisation in the whole bacterial community.

Marine bacterial communities in polar regions are dominated by Alphaproteobacteria, Gammaproteobacteria and Bacteroidetes, with an important contribution of Actinobacteria, Deltaproteobacteria and Archaea (16). The degradation of marine polysaccharides by bacterioplankton requires a high metabolic specialisation, including an appropriate repertoire of enzymes as well as an efficient system to take up the degradation products. Within bacterioplankton, only a few clades have developed the required capacity to efficiently degrade these polymers.

Bacteroidetes is a bacterial phylum specialised in the degradation of polysaccharides (18). Thus, they are a dominant clade in different environments, from soil (19) to the animal intestinal tract (20). In marine environments, Bacteroidetes are more abundant in both polar regions compared to temperate regions (12) for reasons possibly related to substrate availability. The metabolic specialisation of this phylum is given by a genetic repertoire unique to them. Bacteroidetes are known for having a set of carbohydrate-active enzymes (cazymes) that they can secrete to the extracellular medium to degrade complex polymers into simpler, manageable oligomers, which they then take up through specialised transporters and binding proteins. Once in the periplasm, the oligomers can be efficiently degraded with no risk of other bacteria scavenging on the sugars.

In Bacteroidetes species, cazymes and transporters have been found to be clustered in regulated loci known as <u>p</u>olysaccharide <u>u</u>tilisation loci (PULs) (21). This type of locus was first described in the commensal species *Bacteroides thetaiotaomicron* (22).

As mentioned, the Arctic Ocean experiences a bloom of phytoplankton in early summer that constitutes the main primary production event, accompanied by a profusion of polysaccharides. First, the algae growing attached to ice synthesize large amounts of extracellular polysaccharides that are disposed into the water column as the ice melts. And second, the blooming algae themselves produce storage and cell wall polysaccharides that are released because of predation. The composition of the bacterial assemblage changes dramatically during the bloom. Despite previous studies, a community-wide description of the polysaccharide degradation potential of Arctic bacteria is lacking, especially in the context of algal blooms. As most studies have focused on taxonomic descriptions, we still lack functional analyses establishing a connection between enzymatic functions and substrate availability.

In the present work the objective was to describe the bacterial community dynamics using a time-series of metagenomes, and to determine whether the sets of polysaccharide-degrading enzymes of each bacterial group could, at least partially, explain the observed succession of taxa. Metagenomic data contains enough information to describe how different enzymatic functions are distributed among the taxonomic groups, together with their abundance and organisation in the genome.

## Methods

### Sampling

Environmental samples were collected at Dease Strait, lower Northwest Passage, Nunavut, Canada (69.03°N, 105.33°W). Seawater was sampled at 2.5 m deep during spring and summer of 2014 as part of the Ice Covered Ecosystem-CAMbridge Bay Process Study (ICE-CAMPS). Sampling was performed through sampling holes in the ice from 7 March to 24 June. Ice melting started on 8 June, and ice break-up occurred on 19 July. The sampling on 30 July was performed from a boat in ice-free waters.

### DNA extraction and High-Throughput Sequencing

Biomass was collected by filtering approximately 10 L of seawater through series of 20, 3 and 0.22 µm filters. The 0.22 µm filters were kept at −80°C until DNA extraction was performed. Total DNA from filter pieces was extracted using the phenol/chloroform protocol as described elsewhere (23). After thawing, the frozen filters were fragmented into small pieces and then placed into sterile cryovials. Lysis buffer and lysozyme were introduced, resulting in a final lysozyme concentration of 1 mg/ml, followed by a 45-minute incubation at 37°C. Subsequently, sodium dodecyl sulphate (10%) and Proteinase K (0.2 mg/ml) were incorporated into the mixture, and this blend was incubated at 55°C for 1 hour. The lysate underwent two rounds of mixing with an equal volume of phenol/chloroform/isoamyl alcohol (25:24:1, pH 8) and centrifugation at 12,000 × rpm for 10 minutes. To eliminate any residual phenol, a second extraction was performed using chloroform/IAA (24:1). The aqueous phase, containing the DNA, was then concentrated, and purified using Amicon Ultra Centrifugal filters from Millipore. Nanodrop spectrophotometer (NanoDrop 1000 Thermo Fisher) and a Qubit fluorimeter (Thermo Fisher) were used to assess DNA yield and integrity. Finally, the DNA extracts were kept at −80°C until sequencing.

Extracted DNA was sequenced at CNAG (https://www.cnag.crg.eu) utilizing an Illumina HiSeq2000 sequencing platform with a TruSeq paired-end cluster kit, v3. Samples were sequenced in two different batches, the second batch including samples dated May 10, June 1, June 15, and July 30. Details of the sequencing yield are included in Supplementary table 1.

### Bioinformatic analysis

The metagenomic analysis was carried out with the pipeline SqueezeMeta v1.3 (24) and results were later processed with the R package SQMtools (25). This pipeline performed all the necessary steps of metagenomic analysis from raw reads to binning. The pipeline was run in merged mode, which assembled reads from each sample individually using Megahit. Then, the obtained contigs from all samples were combined, de-duplicated and re-assembled. The pipeline then inferred ORFs in the obtained contigs with Prodigal. Taxonomic assignment was done with Diamond homology searches against the NCBI nr database. Functional annotation used a combination of Diamond and HMMer homology searches against PFAM, KEGG and COG databases. Finally, coverage was calculated mapping the reads to the assembly with Bowtie read mapper. Further details on the functioning of the pipeline can be found in Github: https://github.com/jtamames/SqueezeMeta. The results of the obtained assembly are detailed in Supplementary tables 1 and 2. Taxonomic composition was measured in percentages, so changes in these values do not represent actual growth or disappearance of the clades, but just dominance against the others.

### Function Copy-Number Calculation

Copy-number was the chosen measure to estimate the functional content of bacterial clades. It was calculated for taxonomic subsets of the dataset, e.g., considering only sequences annotated as Bacteroidetes. For each of these subsets, copy number was calculated as the ratio between the summed coverage of all genes annotated with a given function and the median coverage of 11 Universal Single-Copy Genes (USiCGs, Supplementary table 3). This transformation noramlises the coverage of a given function relative to the coverage of a set of genes with one single copy expected in each genome. In this way, the obtained copy numbers represented the average number of genes with a given function each clade has.

### Functional Annotation of ORFs

Carbohydrate-active enzymes (cazymes) are enzymes with catalytic activities on carbohydrates. The sequences of all known cazymes have been collected in the CAZy database (26). These enzymes are grouped in five classes, plus an additional class of non-catalytic carbohydrate-binding modules (CBM). The five catalytic classes include glycosyl hydrolases (GH), glycosyl transferases (GT), polysaccharide lyases (PL), carbohydrate esterases (CE) and other accessory redox enzymes (AA).

Functional annotations for the specialised databases CAZy (v.07312020 (27)) and TIGRfam (v4.0 (28)) were added to the PFAM, KEGG and COG annotations that were automatically carried out by SqueezeMeta. Annotations for CAZy were performed with Diamond v2.0.8.146, while annotations against TIGRfam were obtained with HMMer v3.1b2.

As the annotation of enzymatic function by homology is often ambiguous, we developed a consensus annotation (4,29), combining CAZy, PFAM, KEGG and COG annotations to reduce the false-positive rate. To establish an equivalence between PFAM, KEGG and COG identifiers and CAZy families, all sequences in CAZy were annotated in these three databases. The retrieved annotations that were related directly with carbohydrates were selected, removing other more generic domains such as cofactor-binding sites.

We only kept cazymes when at least one of the extra annotations (PFAM, KEGG, COG) was carbohydrate related. This removed spurious cazyme annotations in which a gene resembled a cazyme but was more similar to another protein not included in CAZy, which could only be detected by annotating against broad databases such as PFAM, KEGG and COG. To test this approach, nine reference genomes were retrieved from PATRIC (30). The selected genomes belonged to marine bacterial species that had their curated cazyme annotation published in CAZy (Supplementary table 4). The nine genomes were automatically annotated using the same pipeline used for the metagenomes, and then the CAZy annotation was refined using the proposed consensus approach. The results of this consensus annotation improved overall accuracy with a minimum reduction in recall (Table 1).

**Table 1.** Annotation of different cazyme groups in reference genomes. Rows show the reference curated annotations (Ref.), the automatically retrieved annotations without any filter (Auto.), and those obtained after applying the consensus filter. Columns show the total number of annotations (N) and the recall and accuracy percentages.

|  | GH |  |  | GT |  |  | PL |  |  | CE |  |  |
| --- | --- | --- | --- | --- | --- | --- | --- | --- | --- | --- | --- | --- |
|  | N | Rec. (%) | Acc. (%) | N | Rec. (%) | Acc. (%) | N | Rec. (%) | Acc. (%) | N | Rec. (%) | Acc. (%) |
| Ref. | 458 | - | - | 365 | - | - | 40 | - | - | 30 | - | - |
| Auto. | 1140 | 97 | 39 | 1181 | 100 | 31 | 60 | 93 | 62 | 135 | 97 | 21 |
| Filter | 481 | 94 | 89 | 439 | 99 | 82 | 35 | 80 | 91 | 61 | 97 | 48 |

### PUL Identification

Polysaccharide Utilisation Loci (PULs) were predicted with the criteria used to create the database PULDB (31). A PUL is described as a locus where a *susC-susD* gene pair is present and it is clustered with at least three cazymes. This gene pair comprises a TonB-dependent transporter (*susC*) and a polysaccharide-binding protein (*susD*). Genes were considered to be clustered if they were contiguous in the same direction and less than 102 bp apart, except for cazymes, that could exceed this limit. *SusC* genes were retrieved by looking for associated annotations PF00593 [TonB-dependent receptor], PF07715 [TonB-dependent receptor plug domain] and PF13715 [Carboxy-pepD domain] from the PFAM database, and annotation TIGR04056 [SusC/RagA Family] from TIGRfam; *susD* genes were retrieved with domains PF07980 [SusD Family], PF12741 [SusD and RagB], PF12771[Starch-binding SusD-like] and PF14322 [Starch-binding SusD-like N-terminal] from PFAM (21). Those contigs that had a *susC-susD* tandem and at least 3 cazymes were visualised using the DNA Features Viewer Python library (32) to check the distance and direction criteria.

## Results

### Taxonomic Diversity

The studied time series could be divided in two periods. The first three samples corresponded to late winter, when ice was around 2 m thick and algae had not bloomed, although ice-associated diatom growth has been described in this period. The microalgae population during late winter was dominated by small flagellates. The remaining samples covered the increase in chlorophyll from early May to late June. As ice melting triggered the algal bloom, the chlorophyll peak occurred later (July, Figure 1) compared to temperate regions (March-April).

**Figure 1.**
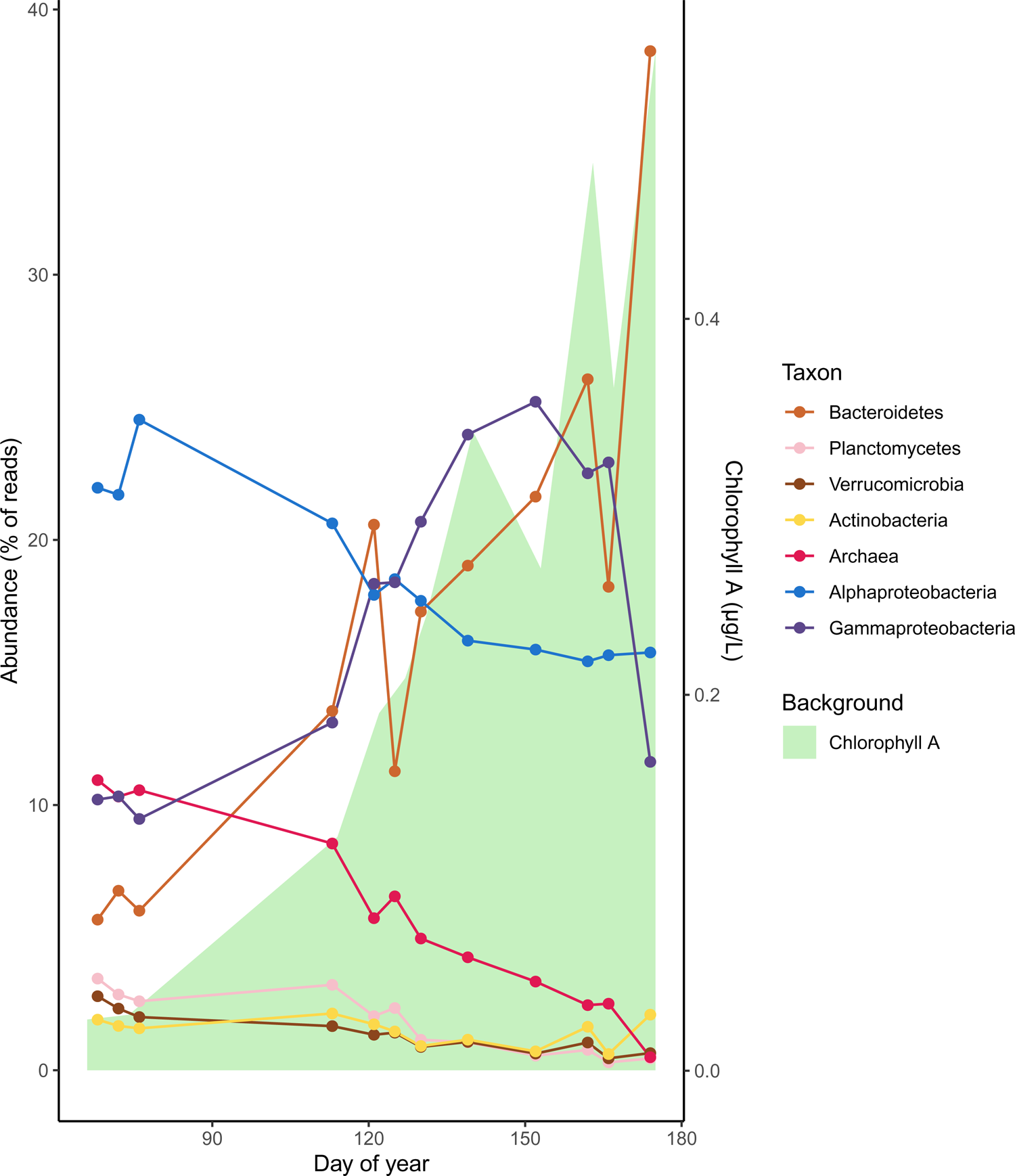
Percentage of reads of different taxonomic groups with time. The background area represents chlorophyll A (µg/L) in water.

In the metagenomic sequences, over 80% of the taxonomically classified reads belonged to phyla Proteobacteria (57.8 %) and Bacteroidetes (24.4%). Within these two phyla, three classes represented over 80% of the reads taxonomically classified at class level: Alphaproteobacteria (33.2%) and Gammaproteobacteria (28.7%) from the phylum Proteobacteria, and Flavobacteriia (21.6%), from the phylum Bacteroidetes.

Alphaproteobacteria was the dominant clade during early spring (23% of reads), with the most abundant annotated orders being Pelagibacterales (50%) and Rhodobacterales (15%). Almost half of the Pelagibacterales reads were classified as *Candidatus* Pelagibacter.

Gammaproteobacteria was the second most abundant class (10% of reads), with a dominance of *Candidatus* Thioglobus (almost 25% of the Gammaproteobacteria reads). Archaea were also abundant during this season, with over 10% of the reads. Bacteroidetes were at that time a minority, contributing roughly over 6% of the reads, with a high proportion of Flavobacteriales (50% of the reads in the phylum). Other minority phyla were Actinobacteria, Verrucomicrobia and Planctomycetes, within the 1.5-3% range each (Figure 1).

As the bloom progressed, Bacteroidetes steadily increased from slightly over 6% of the reads to over 17% in May and over 26% in June. Flavobacteriales were more dominant with time, representing over 70% of Bacteroidetes reads during May and June. *Polaribacter* was the most abundant genus at that time, representing more than 20% of Flavobacteriales reads, despite comprising less than 4% of Flavobacteriales reads in late winter.

Alphaproteobacteria, which were dominant in early spring, decreased to around 17% of the reads with a slight increase of Rhodobacterales, which grew from 15% of the class reads in early spring to over 20% in May and June, reaching a 60% maximum on June 23.

Gammaproteobacteria increased rapidly from 10% to over 20% of the reads, reaching a 25% in early June and decreasing since then back to 11% at the end of the month. The most prominent groups in this class were *Candidatus* Thioglobus (20-30%), Cellvibrionales (5-15%) and the SAR86 cluster (5-10%). The *Cand*. Thioglobus clade (formerly included in the SUP05 clade) peaked at the beginning of the bloom, with almost 30% of the class reads, and decreased there on. Cellvibrionales increased together with the bloom, reaching a maximum of 20% in June and decreasing in the late bloom. The SAR86 clade was most abundant in early spring (10%) and decreased with the bloom, reaching a minimum of 3% in June.

Archaea decreased from 10% of the reads in early spring to around 2% by late June. The reads were mostly assigned to Nitrosopumilales (25-55%). Minority phyla (Actinobacteria, Planctomycetes and Verrucomicrobia), showed a decreasing trend from around 3% in early spring to about 1.5% in late summer.

### Functions in Arctic Communities

Copy number of the most relevant functions for the degradation of polysaccharides revealed that Bacteroidetes, Verrucomicrobia and Planctomycetes had a larger number of degradative enzymes than the rest of the clades (Figure 2). These three phyla had more catalytic cazymes (GH, GT, PL and CE), sulphatases and peptidases than the other clades, though Bacteroidetes stood out as having more glycoside hydrolases.

**Figure 2.**
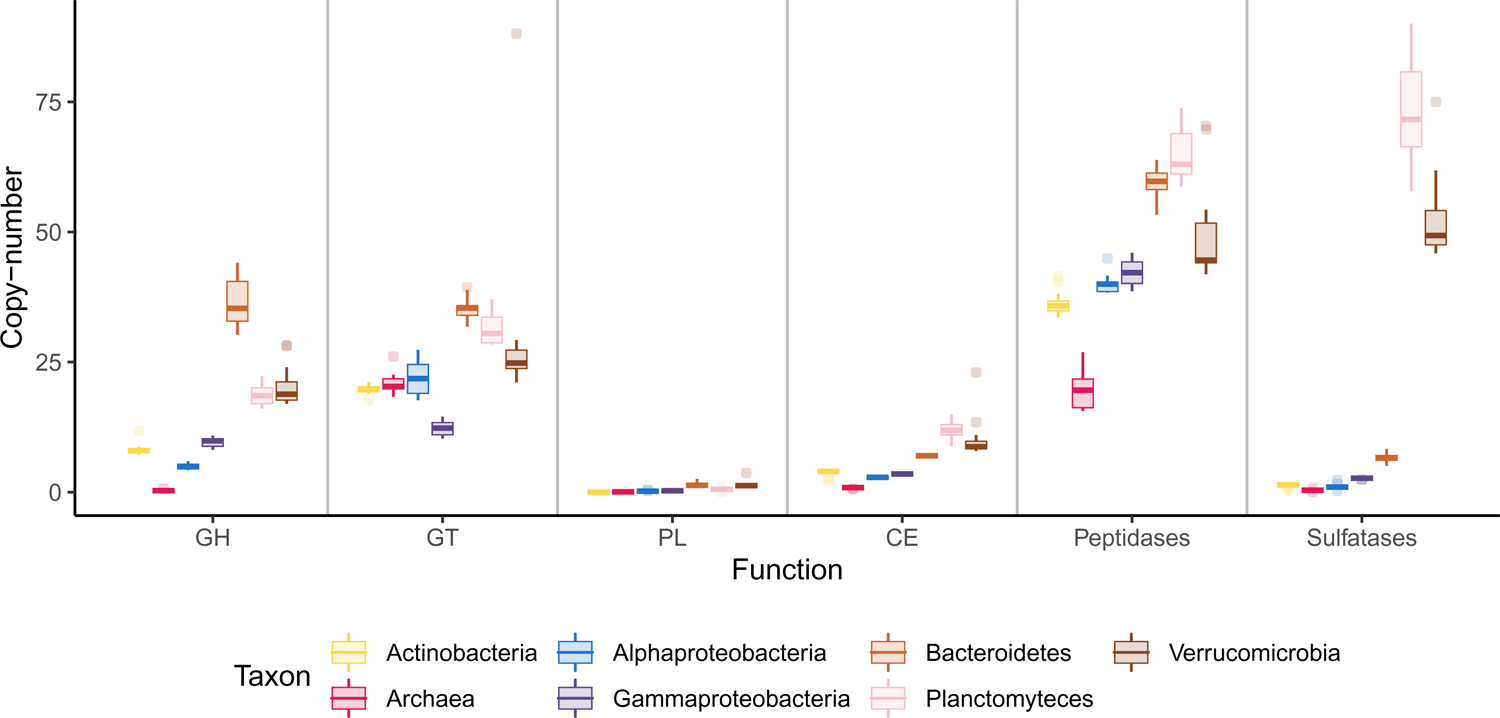
Copy number of relevant functions among taxonomic groups. Each box represents copy number variations across the time series.

Archaea had a singular cazyme profile, having very low copy numbers for most degradative cazymes (GH, PL, CE) and sulphatases, and only having relatively high numbers of GT and peptidases. The main GT families annotated for Archaea were GT1, GT2, GT4 and GT66, related with glycoprotein biosynthesis, showing that these functions are not related with polysaccharide use.

The peptidase:cazyme ratio, which has been used as an indicator of Bacteroidetes species lifestyle, decreased during the bloom (Figure 3) from 1.5 in early spring to almost 1.1 at the point of maximum chlorophyll. Values higher than one have been related to planktonic species, while values closer to or below one are associated with algae-associated species.

**Figure 3.**
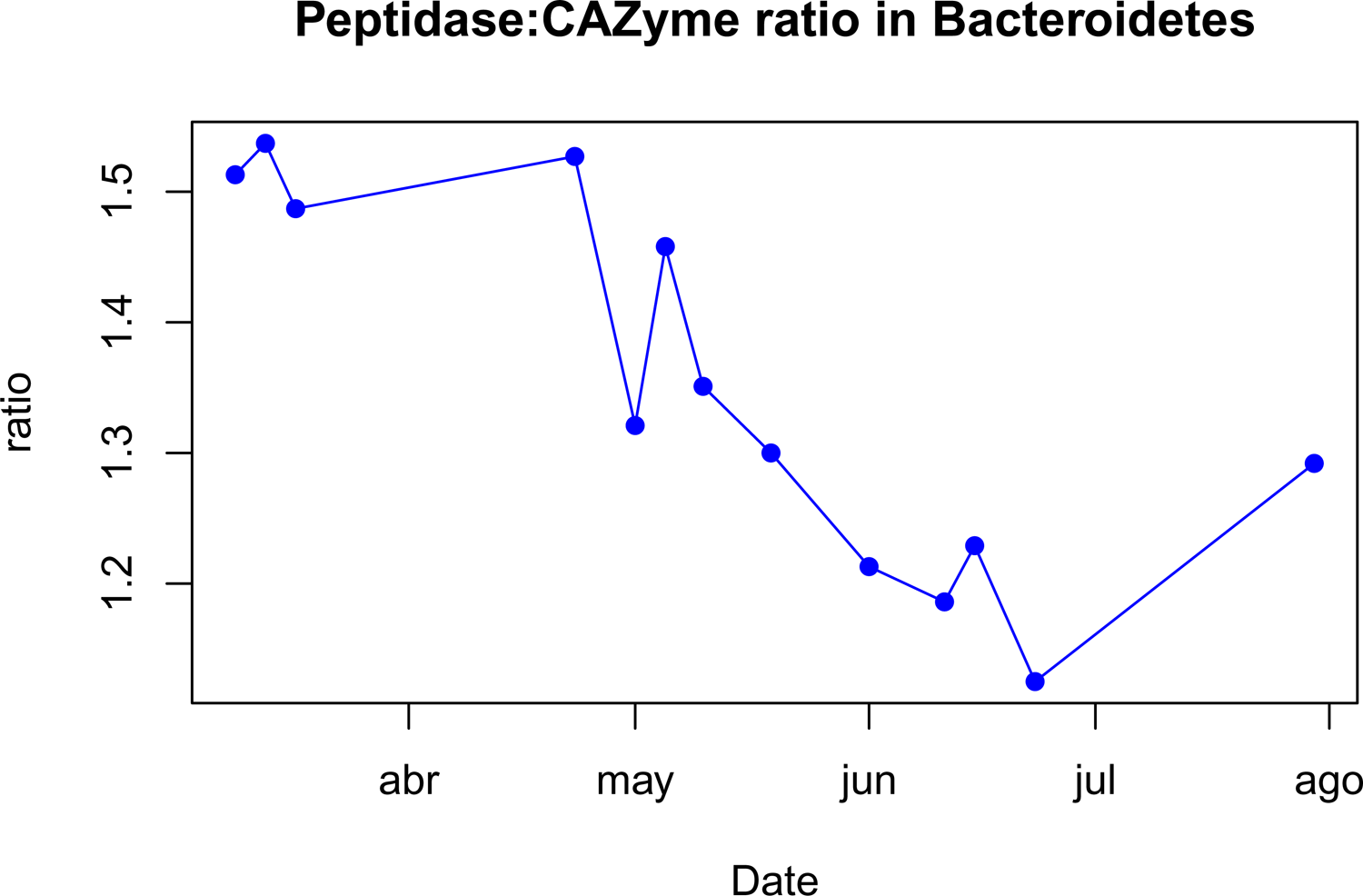
Peptidase:cazyme ratio in Bacteroidetes genomes through the time series.

Sulphatases were distributed unevenly, with extremely high numbers in the Planctomycetes and Verrucomicrobia phyla (73 and 52 copies) and markedly lower numbers in the remaining groups, with Bacteroidetes having at most 7 copies per genome.

### GH Profiles

The distribution of GH families in different taxa shows the potential of each group to degrade different substrates. To compare the enzymatic repertoires of different clades, the Chi-squared distance was calculated for the cazyme profiles of the considered taxa, using proportional copy-number data.

Broad unspecific families such as GH0 (unclassified), GH1, GH2, GH3 or families present in almost all groups (GH13 or GH65) were uninformative. However, most families in CAZy have a narrow functional annotation and are restricted to a few clades.

Some of the GH families that were exclusive or mainly dominated by Bacteroidetes (Supplementary figure 1) were GH30 (glucanase), GH92 (mannosidase), GH130 (mannose phosphorylase) or GH149 (glucose phosphorylase). Other GH families were only shared by Bacteroidetes with Planctomycetes or Verrucomicrobia, such as GH29 (fucosidase) and GH65 (phosphorylase).

Planctomycetes and Verrucomicrobia also had some exclusive hydrolase families, most notably GH33, the only family with sialidase activity found in the dataset, and GH43, a family of arabinases and xylosidases.

Proteobacteria shared two families of lysozymes, GH23 and GH73. Gammaproteobacteria dominated two other families, GH36 (α-galactosidase) and GH42 (β-galactosidase). Alphaproteobacteria had relatively less diversity of enzymes with no clear specialisation (Supplementary figure 1).

Transferases (GT) did not diverge so clearly among the different phyla (Supplementary figure 2). GT profiles diverged the least of all the studied groups, with the lowest Chi-squared distance in GT family composition among the different taxa (Supplementary table 5). Most groups shared the same families, namely GT2, GT4 and GT51, which are involved in peptidoglycan and glycolipid biosynthesis or post-translational modification of proteins, rather than in polysaccharide degradation. GT5 (glycogen synthase) was exclusive to Bacteroidetes, Planctomycetes and Verrucomicrobia and GT66 (glycoprotein biosynthesis), exclusive of Archaea. Proteobacteria shared GT41, involved in the post-translational modifications of adhesins.

Lyases (PL), a group of enzymes that specifically act on uronic acid residues, markedly varied among the groups (Supplementary figure 3). However, most annotated polysaccharide lyases targeted alginate or pectin. Bacteroidetes and Gammaproteobacteria mostly relied on PL6, PL7 and PL17 for alginate lysis. Bacteroidetes had a unique family, PL8, targeting the mannuronic-rich regions in alginate. Planctomycetes and Verrucomicrobia, on the other hand, seemed to rely more on family PL14. Planctomycetes also had a great abundance of PL10, targeting pectin. Verrucomicrobia had less abundance of alginate-targeting lyases, and instead had PL2 and PL10 (pectate lyases) and PL12 (heparin lyase). Annotations in Actinobacteria, Alphaproteobacteria and Archaea were poor due to their low copy numbers.

Most bacteria shared the same families of esterases (CE, Supplementary figure 4), devoted to removing O- and N-decorations from sugar monomers. Most targeted xylan (CE1 to CE7) and murein (CE9 and CE11). CE14 was common in the dataset, but it is poorly characterised in the CAZy database, with no experimental confirmation of the annotated chitobiose deacetylase activity. Verrucomicrobia and Planctomycetes were the only phyla containing CE15, which mainly represents glucuronic acid methylesterases (though no bacterial protein has been characterised). One must be weary of trusting these esterase annotations, as this enzyme group performed poorly in annotation (see Methods).

Differences in the cazyme profiles among taxonomic groups were assessed by calculating the chi-squared distance (Supplementary table 5). This measure is commonly used for proportion data and is suitable to perform metric MDS.

The selected clades diverged the most in GH and PL content, while GT and CE content generated lower distances, indicating more similarity. To better visualise the differences between GH profiles, which are the most abundant and most relevant enzymatic group, a metric MDS ordination was made (Figure 4). In this ordination, Archaea appear clearly apart from the rest of the taxa, showing their highly divergent GH content. Verrucomicrobia and Planctomycetes are represented in proximity, showing that these two phyla have very similar GH profiles, and clearly apart from both Proteobacteria classes, that seem to cluster at the opposite end of the first axis.

**Figure 4.**
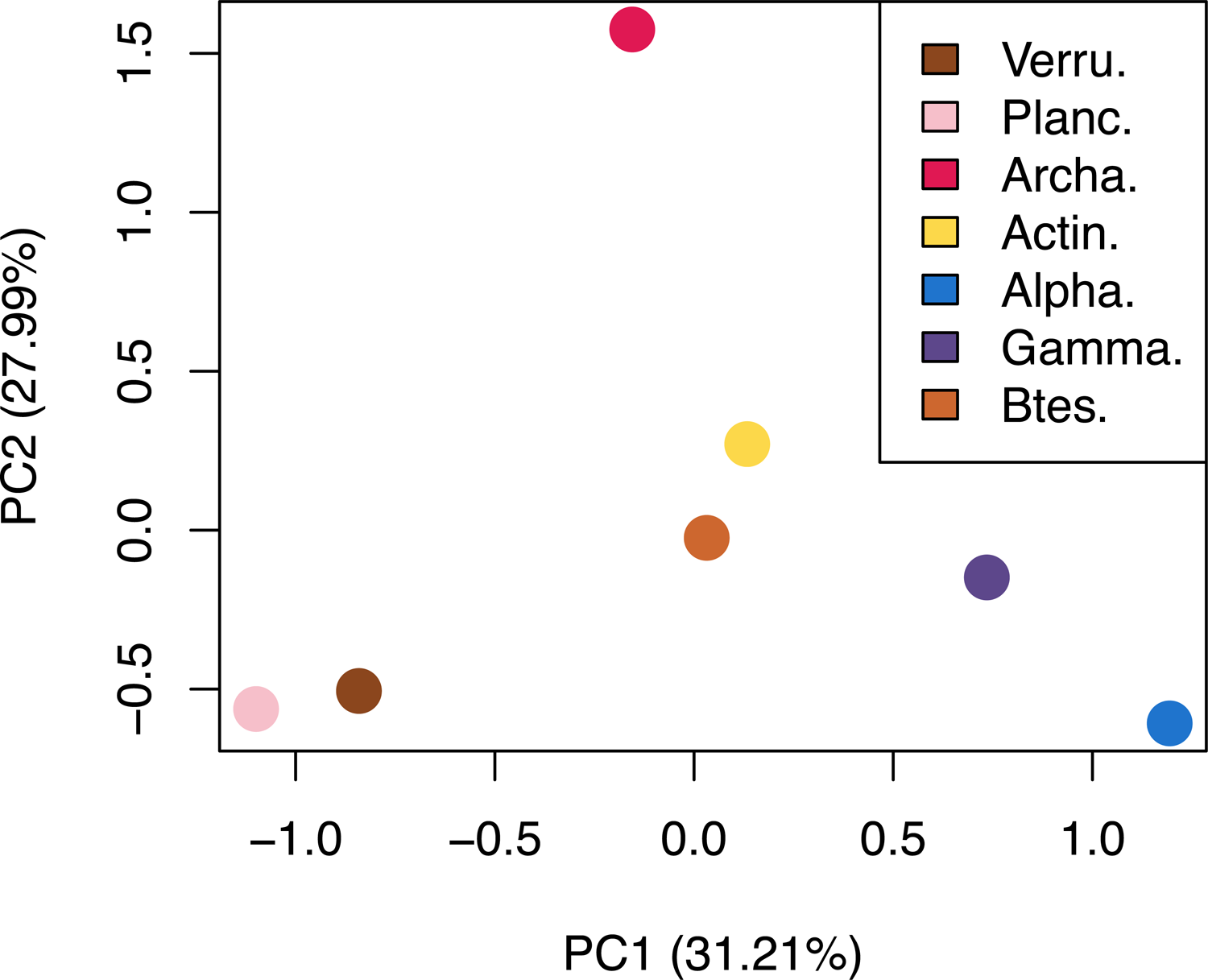
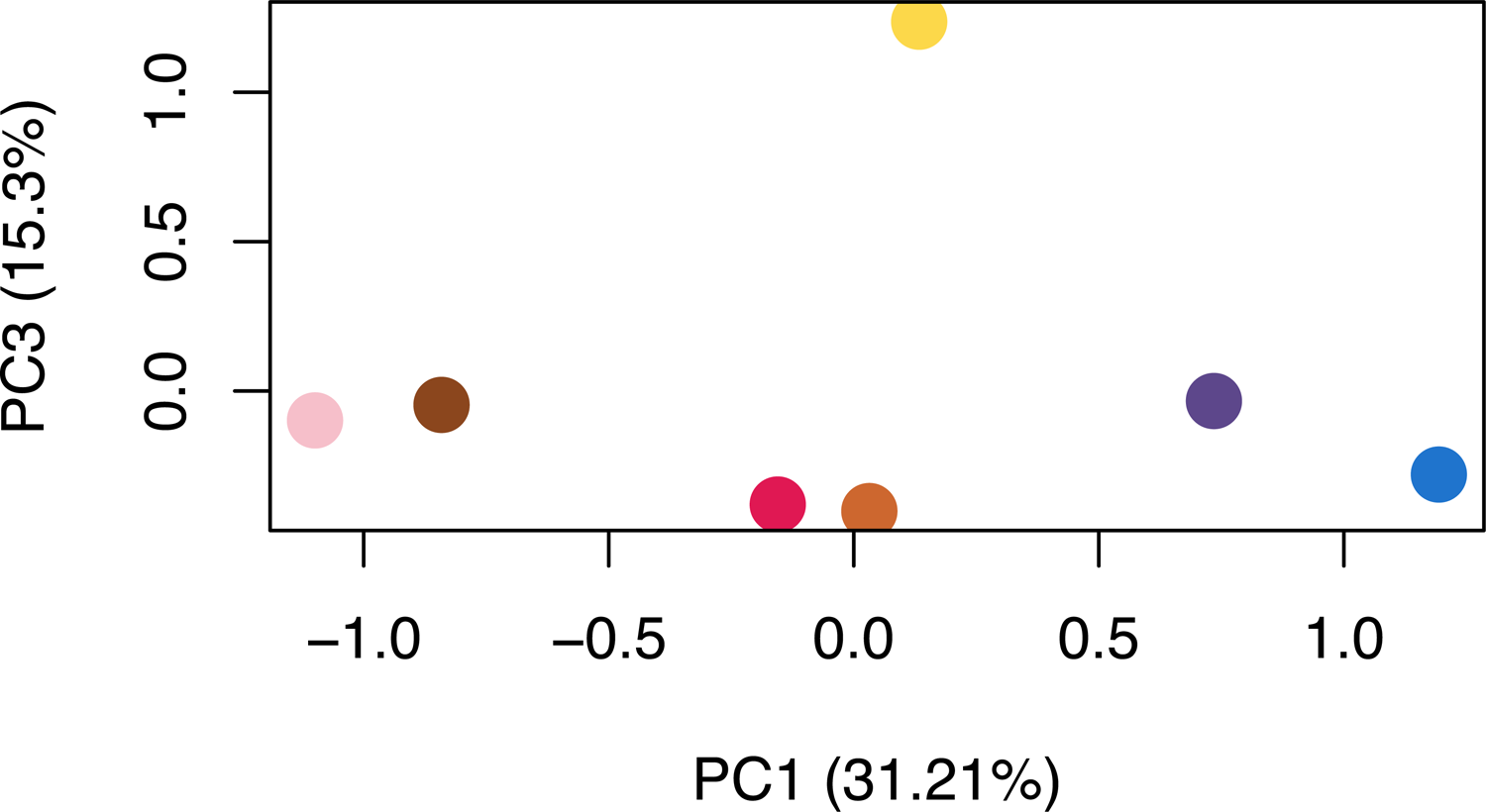
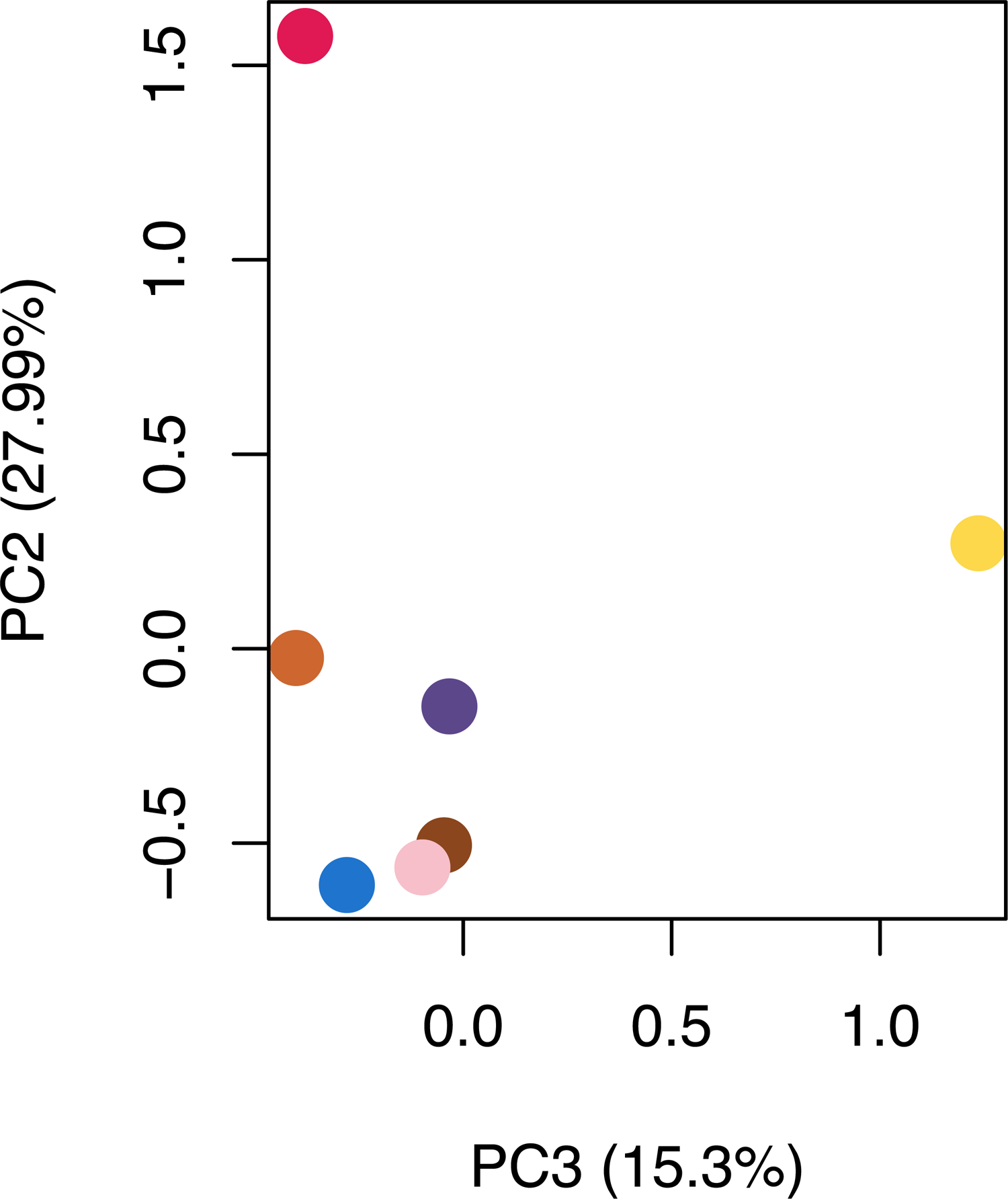
Metric MDS representation of GH profile χ2 distances. From top left to bottom right: PC1 vs. PC2 plot, PC2 vs. PC3 plot, PC1 vs. PC3 plot. Each axis shows the explained percentage of variance. Total explained variance considering the three components: 74. 5%

### Cazymes in Time

The changes in the bacterial community that accompanied the phytoplankton bloom could also be seen in the genetic content of the different clades at the enzyme family level. Bacteroidetes responded drastically to the bloom, with a sharp change in their cazyme content, while the rest of the clades did not show any remarkable change in their cazyme repertoire. The following section will thus focus on the changes at the genomic level in Bacteroidetes.

As spring came about, an increase in the copy number of cazymes could be measured in the Bacteroidetes phylum. Glycosyl hydrolases increased from 30 copies per genome to over 40 in June. Such an increase was neither clearly observed in the rest of the clades nor for any other cazyme group. This increase showed once again that the Bacteroidetes community was shifting towards species with a higher hydrolytic capability as phytoplankton grew.

After seeing that the content of GH in Bacteroidetes increased during the bloom and that this phylum had a high enzyme diversity, the next step was to check if this increase occurred in all GH families equally, or if some functions were more enriched than others during the bloom. For that, the correlation between the copy-number of each GH family and the intensity of chlorophyll was calculated with the Spearman correlation index. P-values for the correlation were corrected using the FDR approach, and only those families with a significant correlation (corrected p-val ≤ 0.01) are reported.

Among the enzymes positively correlated with chlorophyll were those exclusive of Bacteroidetes (Table 2): GH30 (glucanase), GH92 (mannosidase), GH130 (mannose phosphorylase) and GH149 (glucan phosphorylase). Other highly correlated families were GH10 and GH120 (xylanases), GH17, GH81 and GH128 (laminarinases), GH148 (glucomannanase), GH86 (agarase), GH26, GH113 and GH125 (mannosidases) and some glucanase subfamilies in GH5 and GH13 (Supplementary figure 5).

**Table 2.**
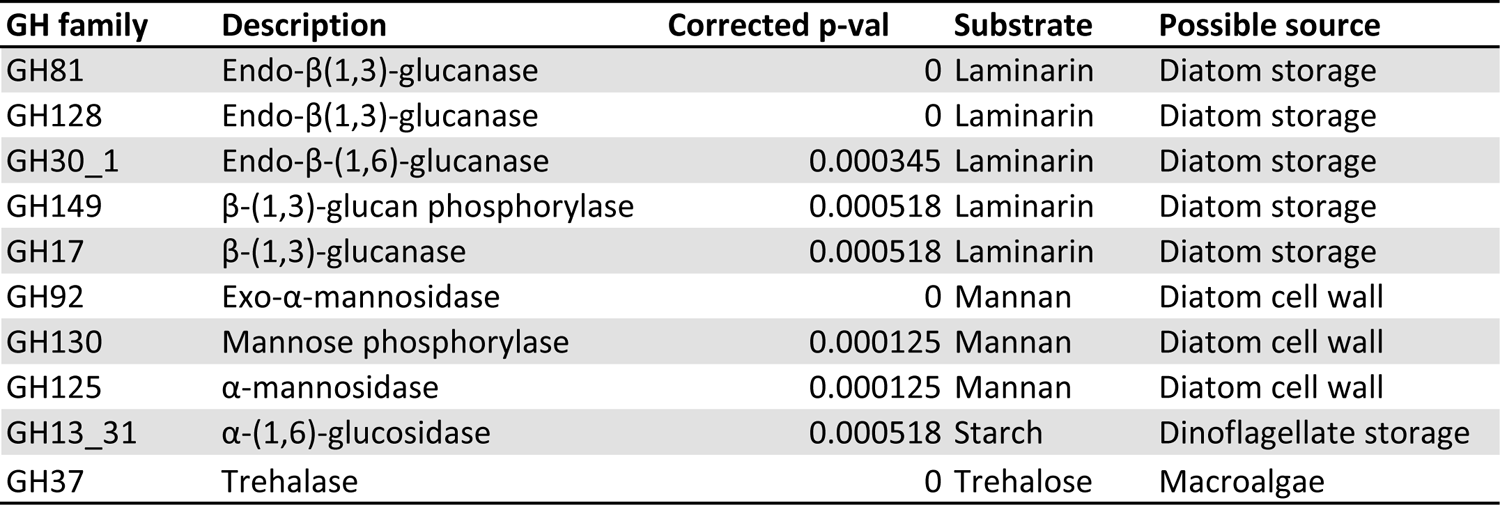
Top 10 GH families whose copy number in Bacteroidetes was positively correlated with chlorophyll with the highest significance. Corrected p-val corresponds to the Spearman’s correlation test after FDR correction.

Out of the 37 GH families and subfamilies that showed a positive correlation, 34 were directly related with polysaccharides, instead of other carbohydrate forms such as disaccharides or glycoconjugates. Two families were related with the utilisation of trehalose (GH37 & GH13_16) and one with sucrose (GH13_18), both anti-freezing and osmo-protectant dissaccharides. Correlated families with the most significant p-values mostly targeted β-(1,3)-glucans (laminarin), β-(1,6)-glucans (laminarin branches) and mannans (Table 2).

Among the families that were negatively correlated with chlorophyll, many were unrelated with algal polysaccharides (Table 3): GH24 and GH25 (lysozymes), GH28 (pectin hydrolase), GH63, GH99 & GH1011 (glycoconjugates); but others represented enzymes that could act on polysaccharides: GH13_36 & GH15 (amylases), GH43 & GH51 (arabinases), GH74 (xyloglucanase), GH95 (fucosidase).

**Table 3.** Top 10 GH families whose copy number in Bacteroidetes was negatively correlated with chlorophyll with the highest significance. Corrected p-val corresponds to the Spearman’s correlation test after FDR correction.

| **GH family** | **Description** | **Corrected p-val** | **Substrate** | **Possible source** |
| --- | --- | --- | --- | --- |
| GH95 | α-L-fucosidase | 0 | Fucoidan | Diatom mucilage |
| GH63 | Glycoglycerate hydrolase | 0 | Glycoglycerate | Cyanobacteria |
| GH99 | Glycoprotein mannosidase | 0 | Glycoproteins | Bacteria |
| GH101 | Endo-α-N-acetylgalactosaminidase | 0.00065 | Glycoproteins | Bacteria |
| GH51 | Hemicellulase | 0.00081 | Hemicellulose | Plants |
| GH25 | Lysozyme | 0.00042 | Murein | Bacteria |
| GH24 | Lysozyme | 0.00076 | Murein | Bacteria |
| GH43_4 | Endo-α-(1,5)-L-arabinanase | 0 | Pectin | Plants |
| GH28 | Polygalacturonase | 0.00065 | Pectin | Plants |
| GH74 | Endo-glucanase specific to xyloglucan | 0 | Xyloglucan | Plants |

### PULs in Bacteroidetes

Differences in the genetic content of the different clades was not restricted to catalytic enzymes. Uptake proteins, namely TonB-dependent transporters (SusC), and associated proteins, such as the polysaccharide-binding protein SusD, also showed a differential distribution among the phyla (Figure 5).

**Figure 5.**
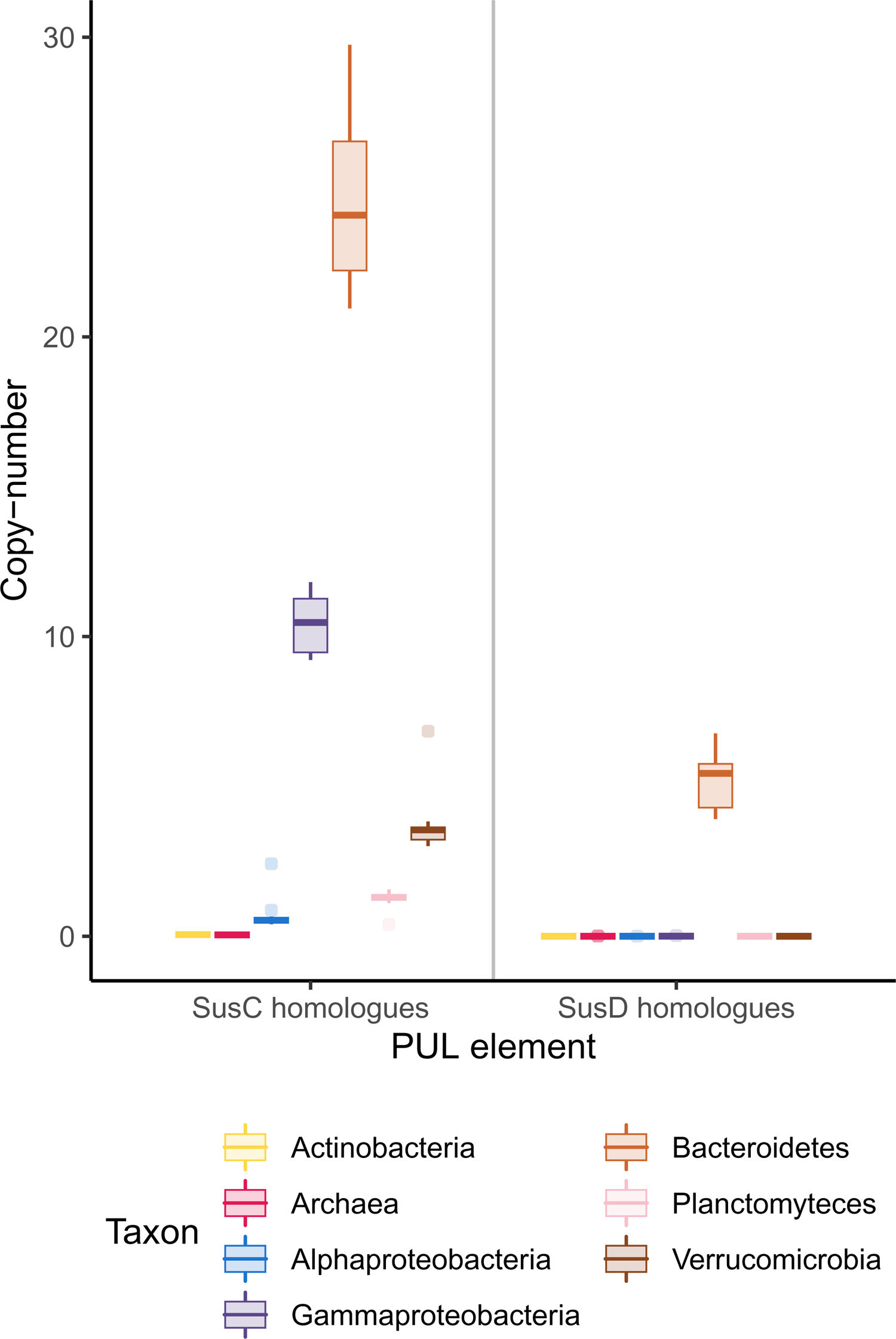
Copy number of SusC and SusD homologues among taxonomic groups. Each box represents copy number variations across the time series.

SusC transporters were most abundant in Bacteroidetes, and the copy number showed a positive correlation with chlorophyll. There were around 21 copies in early spring, rising to almost 27 in June, with a peak of 29 copies in late June (data not shown). Gammaproteobacteria was the second clade with the highest copies of SusC, more than 10, also with a positive correlation with chlorophyll. SusD binding proteins were exclusive of Bacteroidetes, with an average 6 copies per genome.

Some SusD genes were found in Proteobacteria within our dataset, but the low numbers suggest that they might well be bad annotations or an isolated case of gene transfer that does not represent a generalised degradative capability in the clade.

Looking for SusC/D gene tandems, 429 unique tandems were detected in the dataset. Out of these, 400 were in contigs annotated as Bacteroidetes, 26 were unclassified, 2 were annotated as Proteobacteria and 1 as Marinimicrobia.

When adjacent functions were searched to detect a possible PUL around the tandem, about half of the tandems were found to be part of a small contig with only those two genes. On the other hand, 189 tandems were found in contigs big enough to contain at least other three genes, and 61 had at least three cazymes in the contig. Out of these, 34 could be called PULs, as they had three cazymes in proximity and had all nearby cazymes in the same direction as the SusC-D tandem.

Out of these 34 detected PULs, half could be taxonomically classified as down to the Flavobacteriaceae family. These candidate PULs usually included additional genes related with sugar metabolism, such as sulphatases or monosaccharide transporters (MFS, Na-symporters). Some also had a regulatory protein, which is thought to regulate the transcription of the PUL (Figure 6).

**Figure 6.**
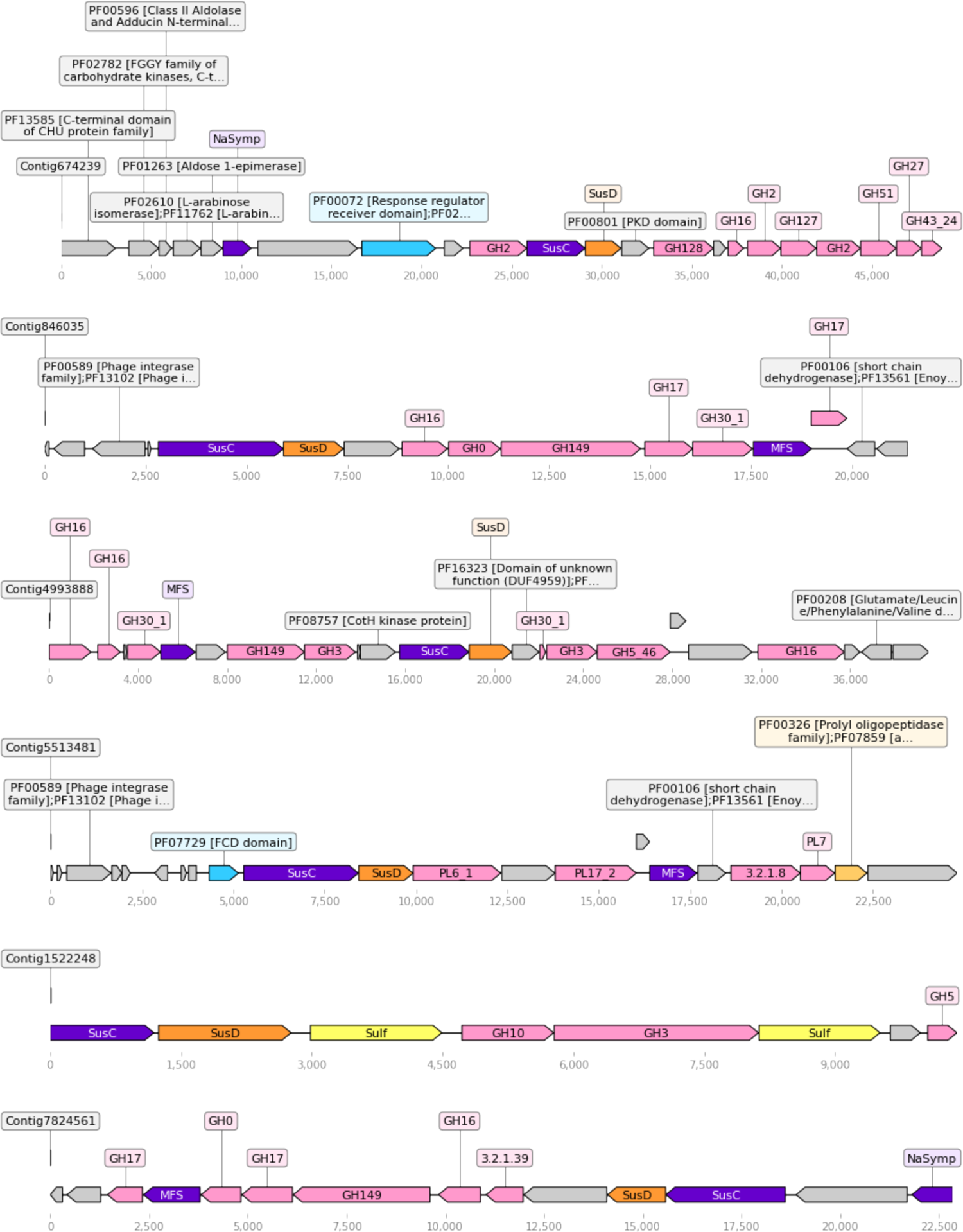
Representation of the biggest PULs found in the dataset. Color key: cazymes (pink), transporters (purple), SusD (orange), sulphatases (yellow), peptidases (gold), regulators (cyan), other (gray). Figure made with dna_feature_viewer.

Within these PULs, functions seem to target a single polysaccharide. As an example, one PUL in contig 846035 was found to contain five enzymes targeting β-1,3-glucans, another in contig 5513482 had three alginate lyases, and one PUL in contig 674239 concentrated up to seven enzymes targeting arabinogalactans.

## Discussion

Algal blooms are seasonal events characterised by a steep increase in phytoplankton abundance in the ocean, a process that fixes tons of carbon from the atmosphere into the biosphere. These blooms trigger the flux of carbon along the trophic chains, with bacterioplankton being the main agent in the remineralisation of the fixed carbon.

Among marine bacteria, Bacteroidetes are a phylum that consistently stands out as taking the most advantage of the new substrates. These bacteria have been studied thoroughly to find that they possess a diverse set of enzymes, known as cazymes, that degrade algal polysaccharides. The current work has extended these studies to Arctic Bacteroidetes, of which little was known.

Bacterial communities in the Arctic show a pre-bloom community dominated by Alphaproteobacteria and Archaea. The Nitrosopumilales clade was the most abundant archaeal group. Within Alphaproteobacteria, the Pelagibacterales clade was dominant during early spring, but was outgrown by Rhodobacterales in the late bloom. These dynamics of Alphaproteobacteria orders has been previously reported in other algal blooms in the North Sea (4). The latter study also showed that this increase in the Rhodobacterales clade continued past the bloom’s peak, which was not covered in the current work. This may indicate a higher capability of the Rhodobacterales clade to use algal polysaccharides.

In the present study, bacterial communities rapidly shifted and Bacteroidetes and Gammaproteobacteria dominated during the bloom. Within Bacteroidetes, Flavobacteriaceae was by far the dominant family, with an important representation of the *Polaribacter* genus. Gammaproteobacteria peaked in the early bloom, decreasing in the late bloom. *Candidatus* Thioglobus, was responsible for most of this early-bloom peak. Cellvibrionales, which have been studied for their plant polysaccharide degradation capabilities, were a minority among the Gammaproteobacteria, but increased together with the bloom.

Within the bacterial community, the polysaccharide degradative capability was distributed unevenly. Bacterial clades significantly diverged in their GH and PL content, but GT and CE profiles were more similar among the studied clades. GT were mostly related with glycoconjugate and bacterial cell wall biosynthesis, not with polysaccharide degradation, which can explain why GT profiles diverged the least.

Archaea seemed to have no capability to degrade complex polysaccharides, and the only functions they contained could be related to glycoprotein synthesis. Proteobacteria had relatively low copy numbers of cazymes, with a higher diversity of functions in Gammaproteobacteria than in Alphaproteobacteria.

Verrucomicrobia and Planctomycetes stood out as having up to tenfold as many sulphatases as the other clades. These high numbers are consistent with those obtained in genomic studies of Planctomycetes (33) and Verrucomicrobia (34) species. These two phyla also had high copy numbers for degradative cazymes (GH, PL and CE) with similar functional profiles. This may indicate some similarities in the strategy and the carbon sources used by Planctomycetes and Verrucomicrobia, which both form part of the PVC superphylum (35).

Verrucomicrobia have been proposed as relevant polysaccharide degraders in Arctic fjords (36), but their low abundance in the Arctic Ocean and the lack of a measurable response to the algal bloom does not suggest such a role in this environment. They have also been reported as potential fucoidan degraders (34), and the presence of GH29 (fucosidase) in their cazyme repertoire supports this. They, together with Placntomycetes, also had a cazyme repertoire with abundant arabinosidases (GH27, GH43), even though arabinose is not one of the main monomers in phytoplankton polysaccharides.

These phyla might well be specialised on lesser-known and hydrolysis-resistant substrates or in minor monomers present as decorations in the main polysaccharides, which may account for their lower abundance as they may feed slowly on these polymers. One of these proposed substrates could be a diatom sulphated fucan polysaccharide that was described as resistant to enzymatic degradation in the North Sea (37). Another possible explanation for this disadvantage could be the absence of siderophore transporters in these phyla, which might hinder their growth by reducing their competitiveness for iron uptake (38).

Although most bibliography comments on the abundance and diversity of cazymes in Bacteroidetes, and they relate their success with that genetic repertoire, the current work shows that other clades, namely Verrucomicrobia and Planctomycetes, have a similar amount and diversity of cazymes, yet they only constitute a minority in Arctic bacterioplankton and do not grow in response to the algal bloom. This shifts the attention of the amount and variety of cazymes to the use that these bacteria make of such enzymes.

As has been reported in other bloom studies, Bacteroidetes was the clade that most rapidly and most intensely grew with the onset of the algal bloom, dominating all the other clades within months.

The increase in the abundance of the phylum was related with species that had a higher cazyme content. Those cazymes related to the degradation of laminarin and mannan were especially enriched, showing a positive correlation with chlorophyll. Laminarin is the main marine storage polysaccharide in diatoms and some studies indicate that diatom cell wall are most likely glucuromannans (7). β-1,4-Mannans have also been detected during phytoplankton blooms and determined to be degraded by bacterial communities (37). Thus, the change in Bacteroidetes species shows a clear trend towards species specialised in diatom polysaccharide degradation.

In this section it is worth analysing the peptidase:cazyme ratio in Bacteroidetes. This ratio has been determined to have a correlation with the lifestyle of different species (39). Bacteroidetes species more adapted to oligotrophic environments tend to have smaller genomes with higher peptidase:cazyme ratios (>1), as is the case of *Polaribacter* MED152 (40), while algae-associated Bacteroidetes feature large cazyme-rich genomes with ratios that can get below 1, as is the case of *Flavobacterium johnsoniae* (41). The decrease in the ratio observed with time suggests that the Bacteroidetes community switched towards species with an algae-associated lifestyle as the algal bloom took place.

The presence of mannan and glucan phosphorylases (GH130, GH149) in this phylum was unexpected, as most marine polysaccharides have sulphate decorations, but no phosphate. A closer look at the CAZy annotation of these families showed that both target oligosaccharides. This may indicate that these enzymes are not secreted to degrade the extracellular polysaccharide, but instead are kept in the intracellular medium or in the periplasm to phosphorylate absorbed oligosaccharides and avoid their diffusion to the extracellular medium.

Bacteroidetes have an exclusive feature that makes them unique among all other clades, the PUL loci. These operons are characterised by two proteins, SusC and SusD. SusD proteins are polysaccharide-binding proteins that, together with SusC transporters, form a complex that binds oligosaccharides and transports them into the periplasm of bacteria. SusD proteins are exclusive to the phylum Bacteroidetes. The abundance and variety of cazymes was not an exclusive feature that could unequivocally explain Bacteroidetes dominance during the bloom. However, these regulated loci can justify the better adaptation of Bacteroidetes to the degradation of algal polysaccharides. The enzymes associated with each SusC/D tandem seemed to be consistent with the idea that each PUL targets a specific polysaccharide (21,42).

The identification of PULs in metagenomic sequences was challenging, as it required the assembly of long contigs to confirm the presence of relevant functions around the SusC/D tandem. This technical problem might be overcome in future studies by complementing the data with long-read sequences.

In future studies, more attention should be placed at the expression of these PULs, as their success might precisely come from a fine-tuned regulation that maximises the benefit obtained by Bacteroidetes when they synthesise the enzymes.

## Aknowledgments

We would like to recognise the in-kind support from the Canadian High Arctic Research Station (CHARS) in the Dease Strain sampling. A. Delaforge assisted greatly in data collection and sample processing. V. Balagué, M. Royo and P. Sánchez from the Marine Bioinformatics Core facility MARBITS at ICM-CSIC Barcelona assisted in initial computational analyses, as well as nucleic acid extractions. Most computation and analyses were carried out at CNB-CSIC.

## Data Availability Statement

Generated sequences were deposited under NCBI BioProject ID PRJNA803814. All custom scripts used for data analysis and other supplementary material used in this work were uploaded to the Github repository https://github.com/redondrio/Arctic_022 for public access.

## Notes

https://www.ncbi.nlm.nih.gov/bioproject/?term=PRJNA803814

https://github.com/redondrio/Arctic_022

### Competing Interest Statement

The authors have declared no competing interest.

