## Supplementary Material for "Specialized Bacteroidetes dominate the Arctic Ocean during marine spring blooms"

**Supplementary information**

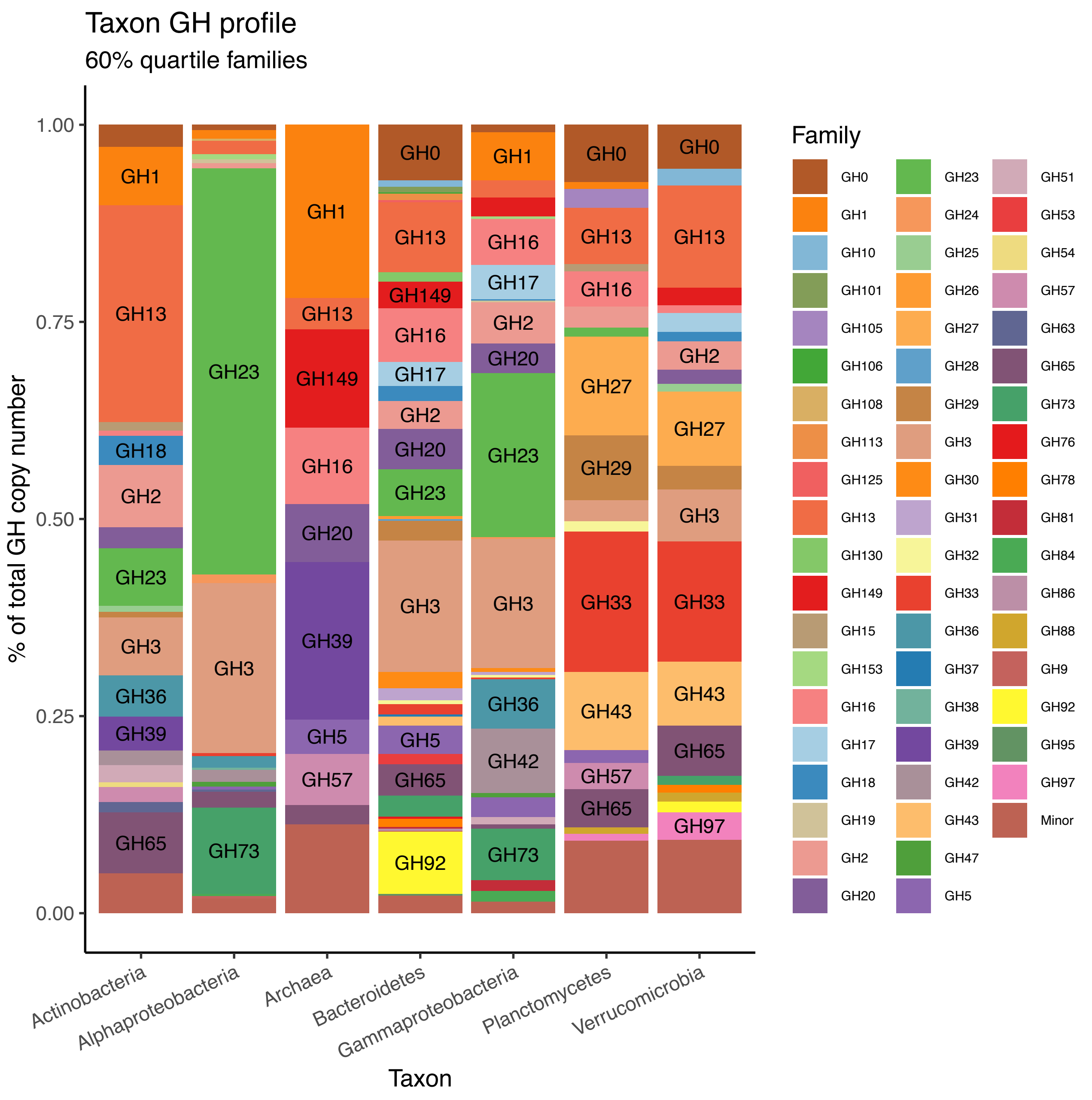


**Supplementary figure 1.** Glycosyl hydrolase (GH) profiles of different taxa, showing the percentage that each family represents out of the total copy number. GH families whose abundance was under the 60% quartile were grouped under the "Minor" tag to ease visualisation.

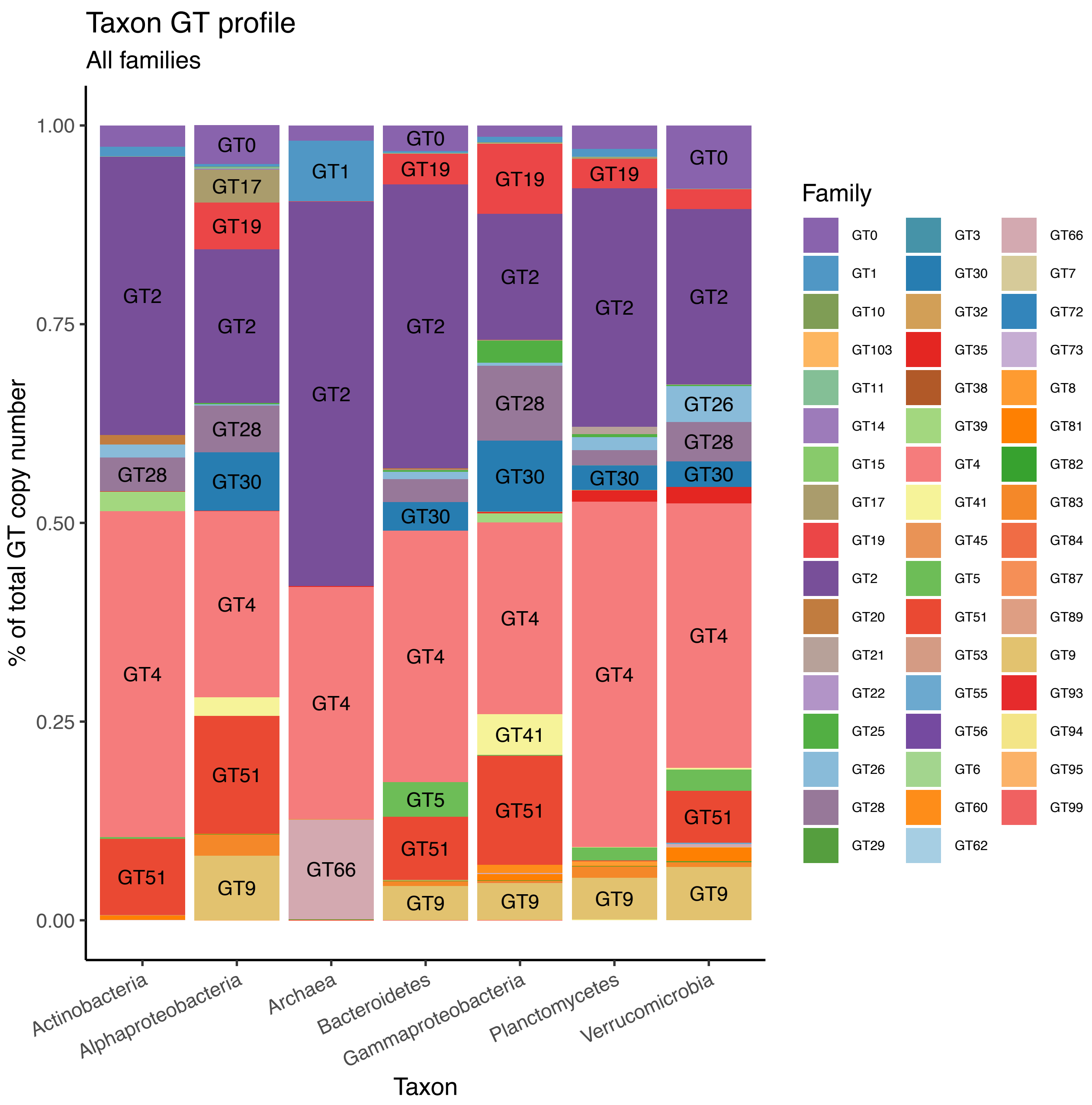


**Supplementary figure 2.** Glycosyl transferase (GT) profiles of different taxa, showing the percentage that each family represents out of the total copy number.

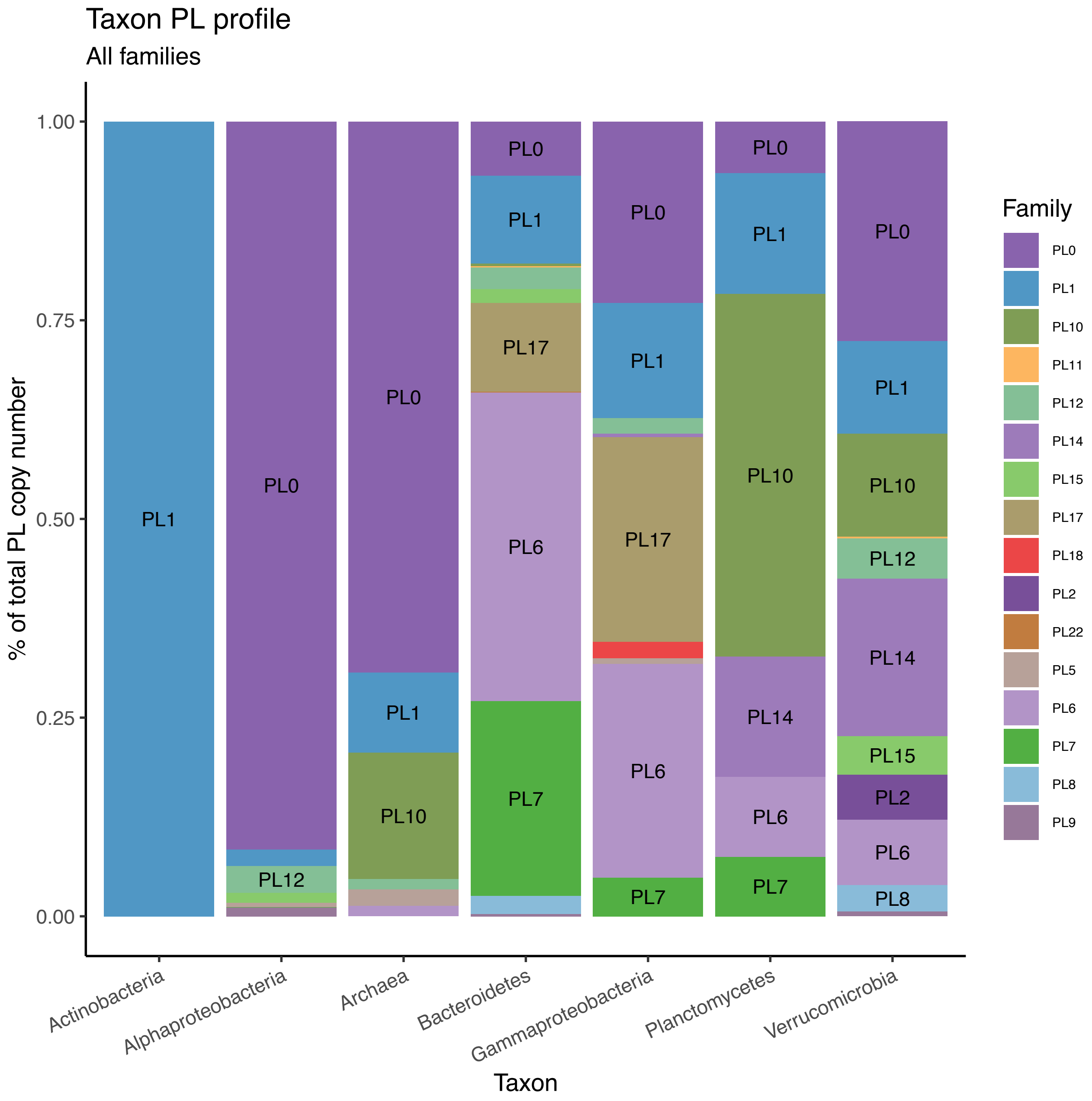


**Supplementary figure 3.** Polysaccharide lyase (PL) profiles of different taxa, showing the percentage that each family represents out of the total copy number.

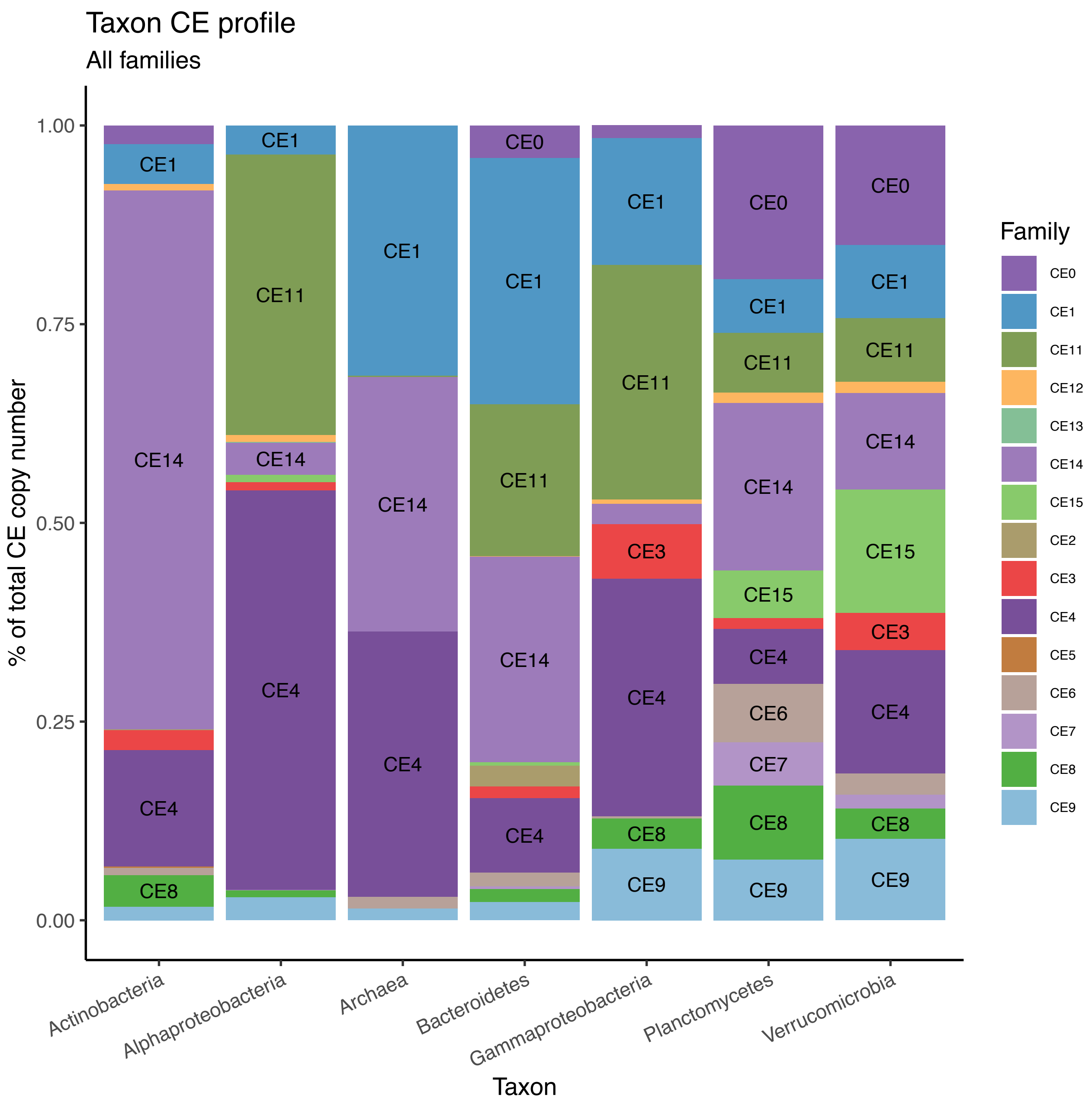


**Supplementary figure 4.** Carboxyl esterase (CE) profiles of different taxa, showing the percentage that each family represents out of the total copy number.

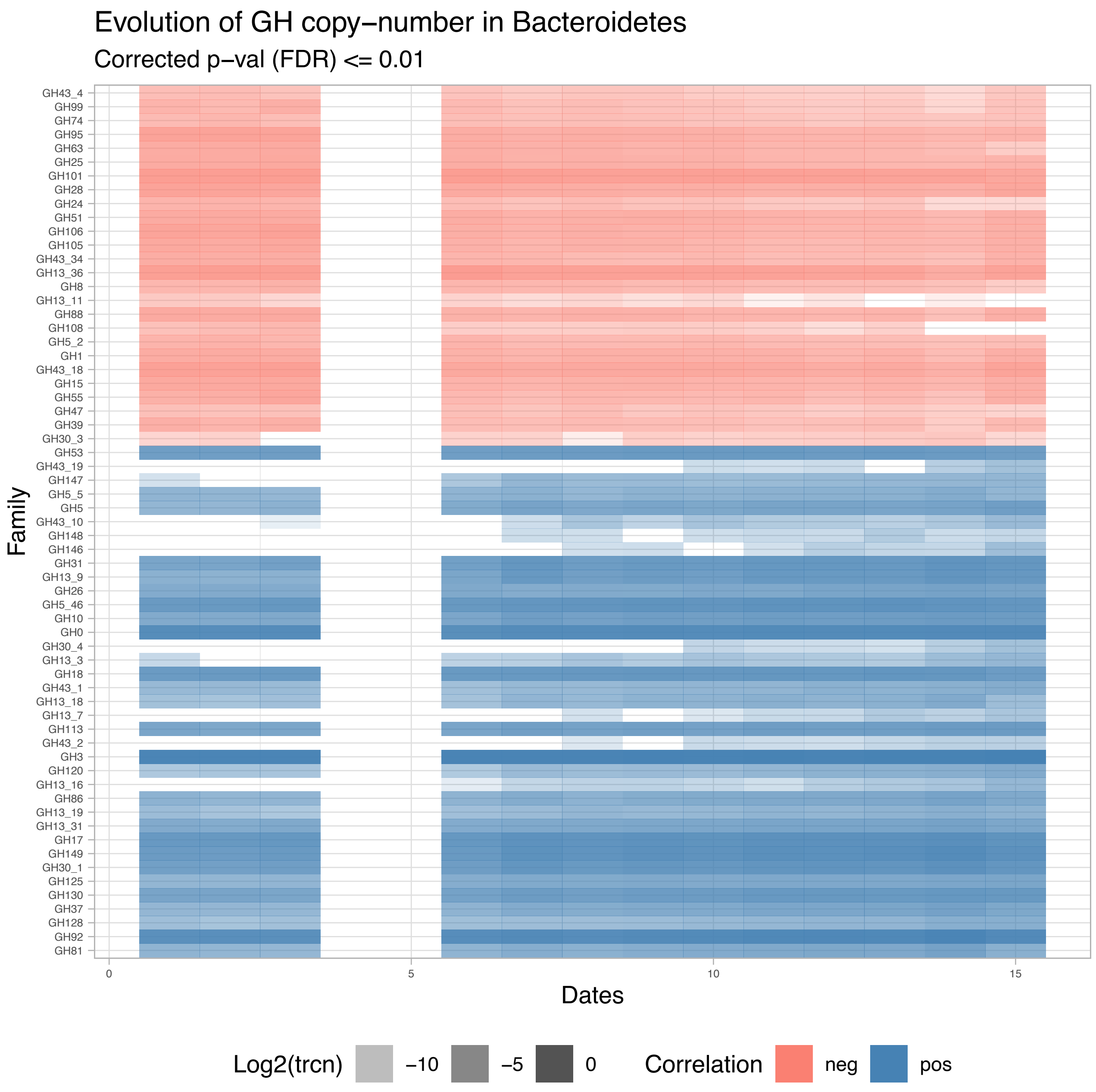


**Supplementary figure 5.** Heatmap showing the copy number of GH families (trcn, log scale) across the time series. Only families showing a significant (corrected p-val ≤ 0.01) correlation with chlorophyll A are shown. Families are ordered by their correlation index, positive (bottom) to negative (top).

| # Sample | Total reads | Reads mapping to assembly | Mapping percent |
|---|---|---|---|
| 9_Mar | 125074094 | 113413500 | 90.7 |
| 13_Mar | 59727336 | 53683468 | 89.9 |
| 17_Mar | 21538100 | 19514486 | 90.6 |
| 23_Apr | 49198506 | 46060030 | 93.6 |
| 1_May | 175932218 | 166272180 | 94.5 |
| 5_May | 138625032 | 126968342 | 91.6 |
| 10_May | 78507842 | 71317694 | 90.8 |
| 19_May | 74655984 | 68633322 | 91.9 |
| 1_Jun | 87152814 | 80861160 | 92.8 |
| 11_Jun | 145220022 | 135669374 | 93.4 |
| 15_Jun | 82445406 | 71402658 | 86.6 |
| 23_Jun | 28946670 | 26775736 | 92.5 |
| 30_Jul | 100424606 | 93968198 | 93.6 |

**Supplementary table 1**. Number of reads and mapped reads for each sample

| **Number of contigs** | **8166358** |
|---|---|
| Total length | 5104467821 |
| Longest contig | 725344 |
| Shortest contig | 200 |
| N50 | 790 |
| N90 | 281 |
| Contigs at superkingdom (k) rank | 6015636 (73.7%), in 4 superkingdoms |
| Contigs at phylum (p) rank | 5330205 (65.3%), in 187 phyla |
| Contigs at class (c) rank | 4210815 (51.6%), in 203 classes |
| Contigs at order (o) rank | 3165768 (38.8%), in 430 orders |
| Contigs at family (f) rank | 2580979 (31.6%), in 696 families |
| Contigs at genus (g) rank | 1413733 (17.3%), in 1654 genera |
| Contigs at species (s) rank | 900102 (11.0%), in 1408 species |
| Number of ORFs | 11057885 |
| Number of rRNAs | 7349 |
| Number of tRNAs/tmRNAs | 50457 |
| ORFs by Aragorn | 50457 |
| ORFs by Prodigal | 10826824 |
| ORFs by barrnap | 7349 |
| ORFs by blastx | 173252 |
| Orphans (no hits) | 2713540 |
| No tax assigned (with hits) | 292959 |
| KEGG annotations | 5380279 |
| COG annotations | 5878147 |
| Pfam annotations | 3349725 |
| CAZy annotations | 537366 |

**Supplementary table 2**. Details of the assembly

| Gene | PFAM annotation |
|---|---|
| RplA | PF00687 [Ribosomal protein L1p/L10e family] |
| RplM | PF00572 [Ribosomal protein L13] |
| RplN | PF00238 [Ribosomal protein L14p/L23e] |
| RplO | PF00828 [Ribosomal proteins 50S-L15, 50S-L18e, 60S-L27A] |
| RplF | PF00347 [Ribosomal protein L6] |
| RpsJ | PF00338 [Ribosomal protein S10p/S20e] |
| RpsK | PF00411 [Ribosomal protein S11] |
| RpsL | PF00164 [Ribosomal protein S12/S23] |
| RpsM | PF00416 [Ribosomal protein S13/S18] |
| RpsG | PF00177 [Ribosomal protein S7p/S5e] |
| RpsH | PF00410 [Ribosomal protein S8] |

**Supplementary table 3.** Universal Single Copy Genes (USiCGs) used for the calculation of copy number and the annotations by which they were retrieved.

| Patric ID | Species |
|---:|---|
| 1347342.6 | Formosa agariphila KMM 3901 |
| 313598.6 | Polaribacter sp. MED152 |
| 313590.8 | Dokdonia sp. MED134 |
| 1798225.3 | Formosa sp. Hel1_31_208 |
| 2058137.3 | Polaribacter sp. ALD11 |
| 1336795.4 | Formosa sp. Hel3_A1_48 |
| 1336804.3 | Polaribacter sp. Hel1_33_78 |
| 1336794.4 | Formosa sp. Hel1_33_131 |
| 376686.1 | Flavobacterium johnsoniae UW101 |

**Supplementary Table 4.** Genomes from Bacteroidetes species used as a reference to test the performance of the consensus annotation filters.

| GH | Verru. | Plancto. | Archaea | Actino. | Alpha. | Gamma. |
|---|---|---|---|---|---|---|
| Plancto. | 1.361 | | | | | |
| Archaea | 2.34 | 2.431 | | | | |
| Actino. | 1.956 | 2.13 | 2.144 | | | |
| Alpha. | 2.256 | 2.408 | 2.596 | 2.14 | | |
| Gamma. | 1.91 | 2.127 | 2.148 | 1.719 | 1.416 | |
| Bacteroidetes | 1.738 | 1.993 | 2.127 | 1.917 | 2.007 | 1.625 |
| GT | | | | | | |
| Plancto. | 0.687 | | | | | |
| Archaea | 1.473 | 1.29 | | | | |
| Actino. | 0.873 | 0.805 | 1.269 | | | |
| Alpha. | 0.99 | 1.037 | 1.65 | 1.131 | | |
| Gamma. | 1.119 | 1.169 | 1.73 | 1.176 | 0.906 | |
| Bacteroidetes | 0.671 | 0.593 | 1.288 | 0.696 | 0.908 | 1.041 |
| PL | | | | | | |
| Plancto. | 2.238 | | | | | |
| Archaea | 1.719 | 2.317 | | | | |
| Actino. | 2.603 | 2.687 | 2.433 | | | |
| Alpha. | 1.95 | 2.529 | 0.97 | 2.697 | | |
| Gamma. | 2.192 | 2.407 | 1.903 | 2.45 | 2.025 | |
| Bacteroidetes | 2.265 | 2.645 | 2.355 | 2.876 | 2.468 | 1.821 |
| CE | | | | | | |
| Plancto. | 0.871 | | | | | |
| Archaea | 1.466 | 1.53 | | | | |
| Actino. | 1.612 | 1.49 | 1.127 | | | |
| Alpha. | 1.544 | 1.726 | 1.374 | 1.795 | | |
| Gamma. | 1.27 | 1.461 | 1.223 | 1.671 | 0.745 | |
| Bacteroidetes | 1.347 | 1.345 | 0.862 | 1.295 | 1.364 | 1.031 |

**Supplementary table 5.** Chi-squared distance values for the four degradative CAZy families (GH, GT, PL and CE) for the seven main taxa. Values across the four CAZY groups are coloured in a white-red scale from lower to higher.
